# Fibroblast-induced mammary epithelial branching depends on fibroblast contractility

**DOI:** 10.1101/2023.03.24.534061

**Authors:** Jakub Sumbal, Silvia Fre, Zuzana Sumbalova Koledova

## Abstract

Epithelial branching morphogenesis is an essential process in living organisms, through which organ-specific epithelial shapes are created. Interactions between epithelial cells and their stromal microenvironment instruct branching morphogenesis but remain incompletely understood. Here we employed fibroblast-organoid or fibroblast-spheroid co-culture systems and time-lapse imaging to reveal that physical contact between fibroblasts and epithelial cells and fibroblast contractility are required to induce mammary epithelial branching. Pharmacological inhibition of ROCK or non-muscle myosin II, or fibroblast-specific knock-out of *Myh9* abrogate fibroblast-induced epithelial branching. Furthermore, fibroblast-induced branching requires epithelial proliferation and is associated with distinctive epithelial patterning of YAP and ERK activity along organoid branches, which is dependent on fibroblast contractility. Together, we identify fibroblast contractility as a novel stromal factor driving mammary epithelial morphogenesis. Our study contributes to comprehensive understanding of overlapping but divergent employment of mechanically active fibroblasts in developmental versus tumorigenic programs.

## Introduction

Efficient formation of large epithelial surfaces in limited organ volumes is achieved through branching morphogenesis (Affolter et al., 2009). The underlying processes of epithelial morphogenesis, including epithelial cell proliferation, migration, intercalation, differentiation, and death, are regulated by both internal genetic programs as well as external cues provided by systemic signals (such as hormones) and local organ-specific microenvironment (Affolter et al., 2009; Goodwin and Nelson, 2020; Wang et al., 2017). The mammary gland is the ideal tissue paradigm for stochastically branching epithelia. Mammary morphogenesis starts in the embryo, but the majority of branch bifurcations and ductal elongation takes place postnatally during puberty. The microenvironment of the mammary epithelium is a dynamic entity that consists of extracellular matrix (ECM) and stromal cells, including fibroblasts. Fibroblasts lay adjacent to the epithelium and have been well recognized as master regulators of mammary epithelial morphogenesis during puberty through production of growth factors (Koledova et al., 2016; Kouros-Mehr and Werb, 2006; Sumbal and Koledova, 2019; Wiseman and Werb, 2002; Zhao et al., 2017) and ECM molecules (Brownfield et al., 2013; Hammer et al., 2017; Jones et al., 2019; Koledova et al., 2016; Nerger et al., 2021; Peuhu et al., 2017; Sumbal and Koledova, 2019; Zhao et al., 2017) necessary for mammary epithelial growth and branching (Sumbal et al., 2020). However, the dynamics of the epithelial-fibroblast interactions during mammary branching morphogenesis as well as whether fibroblasts contribute to shaping of mammary epithelium through additional mechanisms have remained unknown.

Microenvironment of several developing organs has been shown to govern epithelial patterning by dynamic cues of mechanically active cells. Dermal cells in chick skin determine feather buds by mechanical contraction (Shyer et al., 2017), intestinal vilification is dependent on compression by smooth muscle cells (Shyer et al., 2013), and embryonic lung mesenchyme promotes epithelial bifurcation by mechanical forces (Goodwin et al., 2019; Kim et al., 2015; Palmer et al., 2021). However, it has not been elucidated whether the mammary microenvironment contains an instructive component of mechanically active cells as well.

To answer this question, we performed live imaging and functional analysis of co-cultures of primary mammary epithelial organoids with primary mammary fibroblasts. Analogously to primary mammary organoids treated with fibroblast growth factor 2 (FGF2), a well-established model of mammary branching morphogenesis driven by paracrine signals (Ewald et al., 2008), our *in vitro* co-culture model provides a unique window into fibroblast-epithelial interactions during pubertal mammary branching morphogenesis. It enables visualization of stromal fibroblasts during dynamic morphogenetic processes, which are otherwise largely inaccessible *in vivo* due to light-scattering properties of mammary adipose tissue. In this work, we show that physical contact between fibroblasts and epithelial cells, and actomyosin-dependent contractility of fibroblasts are required for branching morphogenesis. Importantly, we demonstrate successful reconstitution of budding morphogenesis by 3D co-culture of contractile fibroblasts in breast cancer spheroids that normally do not form buds. Our results reveal a novel role of fibroblast contractility in driving epithelial branching morphogenesis.

## Results

### Fibroblast-induced branching of organoids does not reproduce FGF2-induced budding

To uncover the role of fibroblasts in epithelial morphogenesis, we investigated differences between organoid budding induced solely by paracrine factors (using primary mammary organoids exposed to exogenous FGF2) and organoid branching induced by fibroblasts (using organoids co-cultured with primary mammary fibroblasts in the absence of any exogenous growth factor). Addition of FGF2 or fibroblasts to mammary organoid cultures both induced branching of epithelial organoids (Figure 1A, B, Suppl. Video 1) but examination of the resulting organoid morphogenesis revealed important differences in dynamics and epithelial architecture in the two conditions. First, organoids co-cultured with fibroblasts developed bigger but less numerous branches (Figure 1A, C). Second, they branched half-day to one day earlier than organoids treated with FGF2 (Figure 1A, D) and the branches were developed rapidly, including the development of negative curvature at the root of the branch (Figure 1E). Third, while FGF2-induced epithelial branching involved epithelial stratification as previously reported (Ewald et al., 2008), co-culture with fibroblasts did not perturb the epithelial bilayer with its lumen (Figure 1A, G). These results suggest that fibroblasts and exogenous FGF2 drive organoid branching by different mechanisms. While the mechanism of FGF2-induced organoid budding was previously described in detail to begin with epithelial proliferation and stratification (Huebner et al., 2014) followed by budding and ERK-dependent and proliferation-independent bud elongation (Huebner et al., 2016), how fibroblasts induce organoid branching remains unanswered (Figure 1F).

**Figure 1.**
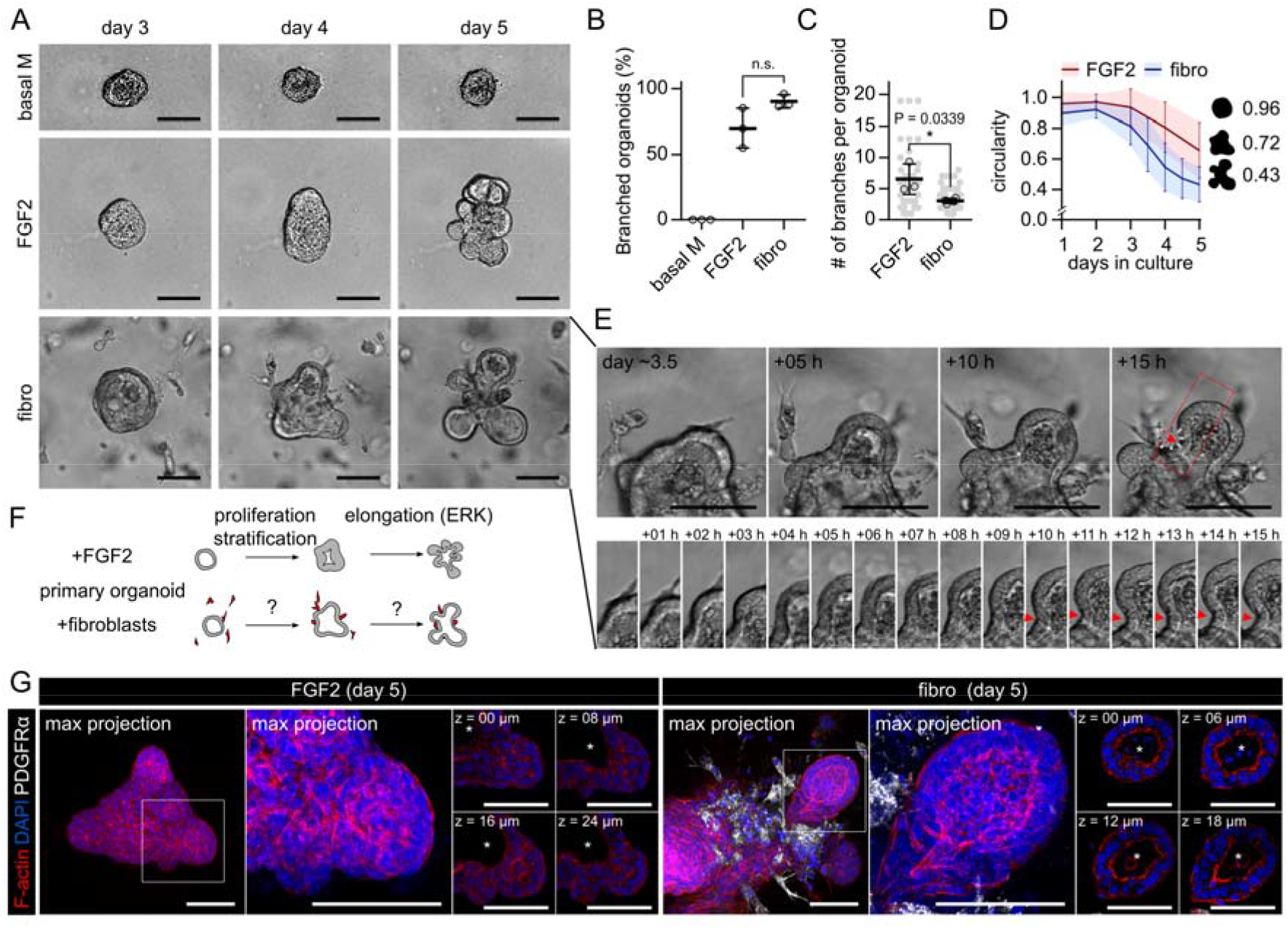
Fibroblast-induced branching of organoids does not reproduce FGF2-induced budding. **A.** Snapshots from time-lapse imaging of primary mammary organoids in basal organoid medium (basal M) without any FGF supplementation (top), basal M with FGF2 (middle), or co-cultured with primary mammary fibroblasts (fibro) in basal M with no FGF supplementation (bottom). Scale bar: 100 μm. Full videos are presented in Suppl. Video 1. **B**. Quantification of percentage of branched organoids per all organoids in the conditions from A. The plot shows mean ± SD, each dot represents biologically independent experiment, n = 3. Statistical analysis: Two-tailored t-test. **C**. Quantification of number of branches per branched organoids in conditions from A. The plot shows mean ± SD, each lined dot shows mean from each experiment, each faint dot shows single organoid measurement, n = 3 biologically independent experiments, N = 20 organoids per experiment. Statistical analysis: Two-tailored t-test. **D.** Quantification of organoid circularity in conditions from **A**, the lines represent mean, the shadows and error bars represent ± SD, n = 3 biologically independent experiments, N = 20 organoids per experiment. The schemes show representative shape of given circularity. **E**. Detailed images of branch development in co-culture with fibroblasts from A. Scale bar: 20 μm. **F**. A scheme depicting differences between organoid budding induced by exogenous FGF2 and organoid branching in co-culture with fibroblasts. **G**. Maximum intensity projection of F-actin (red), DAPI (blue) and PDGFRα (white) in organoid with exogenous FGF2 or with fibroblasts (fibro). Zoom-in area from the box is depicted as maximum projection and single z slices. Scale bar: 100 μm.

### Endogenous paracrine signals are not sufficient to induce organoid branching in co-cultures

The ability of exogenous FGF2 to promote organoid budding (Ewald et al., 2008) (Figure 1A) well demonstrates the importance of paracrine signals for epithelial branching, although FGF2 amount used in *in vitro* branching assays likely exceeds physiological values *in vivo*. Therefore we sought to determine, whether endogenous FGFs or other paracrine signals produced by fibroblasts in co-cultures (Koledova et al., 2016, 1; Sumbal and Koledova, 2019) are sufficient to drive organoid branching. First to test the involvement of FGF signaling, we inhibited either FGF receptors (FGFRs) using SU5402, or ERK, a common signaling node of all receptor tyrosine kinases using U0126. As expected, both inhibitors abolished branching induced by exogenous FGF2 (Figure 2A, B). However, in the co-cultures with fibroblasts, the same concentration of inhibitors did not abolish branching, albeit slightly reduced organoid growth (Figure 2A, B), suggesting that paracrine signaling via FGFR-ERK pathway is not the only mechanism driving organoid branching.

**Figure 2.**
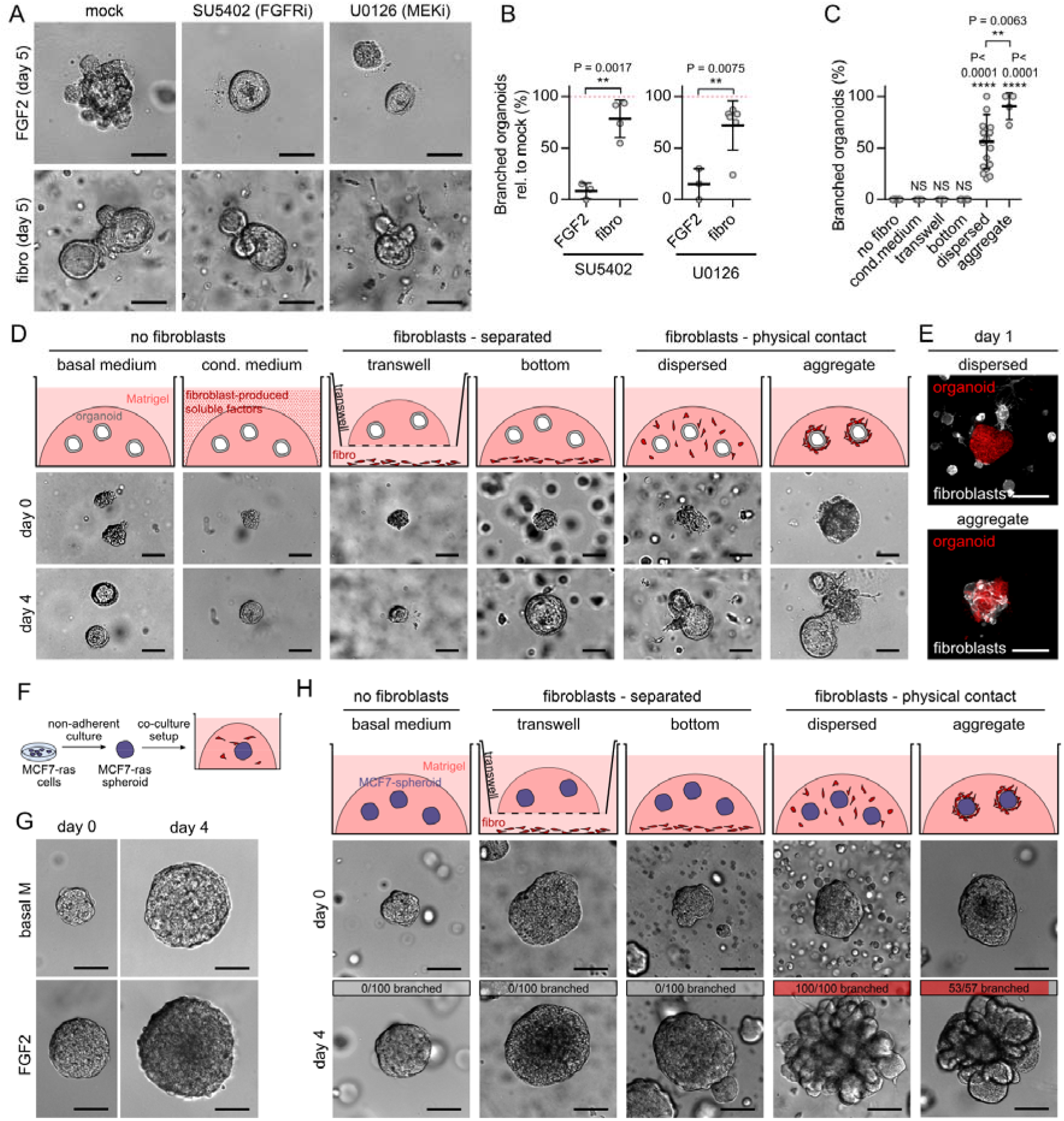
Endogenous paracrine signals are not sufficient to induce organoid branching. **A.** Representative organoids cultured with exogenous FGF2 or with fibroblasts (fibro), treated with inhibitor of FGFR (SU5402) and MEK (U0126). Scale bar: 100 μm. **B**. Quantification of branched organoid per all organoids, relative to mock. The plots show mean ± SD, each dot represents biologically independent experiment, n = 3-5, N = 20 organoids per experiment. Statistical analysis: Two-tailored t-test. **C**. Quantification of organoid branching in different co-culture set-ups. The plot shows mean ± SD, each dot represents biologically independent experiment, n = 16 independent experiments for “no fibro”, 5 for “cond. medium”, 5 for “transwell”, 12 for “bottom”, 16 for “dispersed”, and 4 for “aggregate”, N = 20 organoids per experiment. Statistical analysis: Multiple *t-* tests compared to control “no fibro”, or indicated by the line. **D**. Schemes and images on day 0 and day 4 of different organoid-fibroblast co-culture set-ups. Scale bar: 100 μm. **E**. Dispersed and aggregated co-culture of LifeAct-GFP fibroblast (white) and tdTomato organoid (red) at the beginning of the culture. Scale bar: 100 μm. **F**. A scheme of MCF7-ras spheroid co-culture setup. **G.** Representative MCF7-ras spheroid cultured in basal organoid medium (basal M) or basal M with exogenous FGF2. Scale bar: 100 μm. **H**. Schemes and images on day 0 and day 4 of different spheroid-fibroblast co-culture set-ups. Top grey and red bars indicate proportion of branched spheroids out of all spheroids per condition, n = 3-5 independent experiment, N = 20 spheroids per experiment. Scale bar: 100 μm.

To probe the involvement of other paracrine signaling in fibroblast-induced organoid branching, namely to test if fibroblast paracrine signaling alone is sufficient to induce organoid branching, or if other mechanisms involving fibroblast-epithelial proximity or contact are involved, we performed an array of different types of organoid (co-)culture set-ups (Figure 2D, top). When we provided unidirectional fibroblasts-to-epithelium paracrine signals by culture of organoids in fibroblast-conditioned medium, no organoid branching was observed (Figure 2C, D). When we allowed bidirectional paracrine signals by co-culture of organoids with fibroblasts in the same well but the organoids and fibroblasts were separated by a transwell membrane or by a thick layer of Matrigel, no organoid branching was observed either (Figure 2C, D). However, when we allowed both paracrine signals and physical contact between organoid and fibroblasts by co-culturing them together either dispersed in Matrigel or as aggregates of fibroblasts on top of organoids embedded in Matrigel, we observed organoid branching (Figure 2C-E). These results demonstrated an essential requirement of fibroblast-epithelium contact for fibroblast-induced organoid branching thus revealing that fibroblast-secreted paracrine factors are not sufficient to initiate branching.

### MCF7-ras spheroids recapitulate fibroblast-induced branching of organoids

To further test the requirement of fibroblast-epithelium contact for fibroblast-induced epithelial branching, we developed a simpler co-culture system, where mammary fibroblasts were co-cultured with MCF7-ras breast cancer cell line spheroids (Figure 2F) instead of organoids from normal mammary epithelium. The advantage of MCF7-ras spheroids is that the spheroids grow constantly due to constitutively active RAS kinase and unlike normal epithelium they do not respond to exogenous FGF2 (Figure 2G) or EGF (Suppl. Figure 1A) by branching. We found that similarly to normal epithelium, MCF7-ras spheroids remained round in fibroblast co-cultures, which did not allow physical contact with fibroblasts, but developed numerous buds when physical contact with fibroblasts was allowed (Figure 2H; Suppl. Figure 1B). These results demonstrated that fibroblasts are able to promote epithelial budding even in a system that is morphogenetically unresponsive to paracrine signals.

### Fibroblasts form physical contact with organoids

To gain more insights into the mechanism of fibroblast-induced organoid branching, we examined organoid branching in the co-cultures in higher resolution. First, we performed a co-culture experiment with organoids labeled by mTomato and GFP-tagged fibroblasts. The time-lapse movies unveiled that fibroblasts came in close contact with the epithelium early during the co-culture and remained there during branching (Figure 3A, Suppl. Video 2). On the branched organoids, fibroblasts were exclusively located around the necks of the nascent branches and sat directly in contact with the epithelium (Figure 3B). Immunofluorescence staining of epithelial markers revealed that fibroblasts formed contacts with KRT5 positive myoepithelial cells (Figure 3C, Suppl. Video 3). Transmission electron microscopy of the co-cultures confirmed the close proximity between the fibroblasts and the epithelium, with a thin layer of ECM in between (Figure 3D). Using immunostaining we detected laminin α5, a basal membrane component, between organoid and adjacent fibroblast (Figure 3E, F). These data suggest that fibroblasts form contacts with epithelium via ECM.

**Figure 3.**
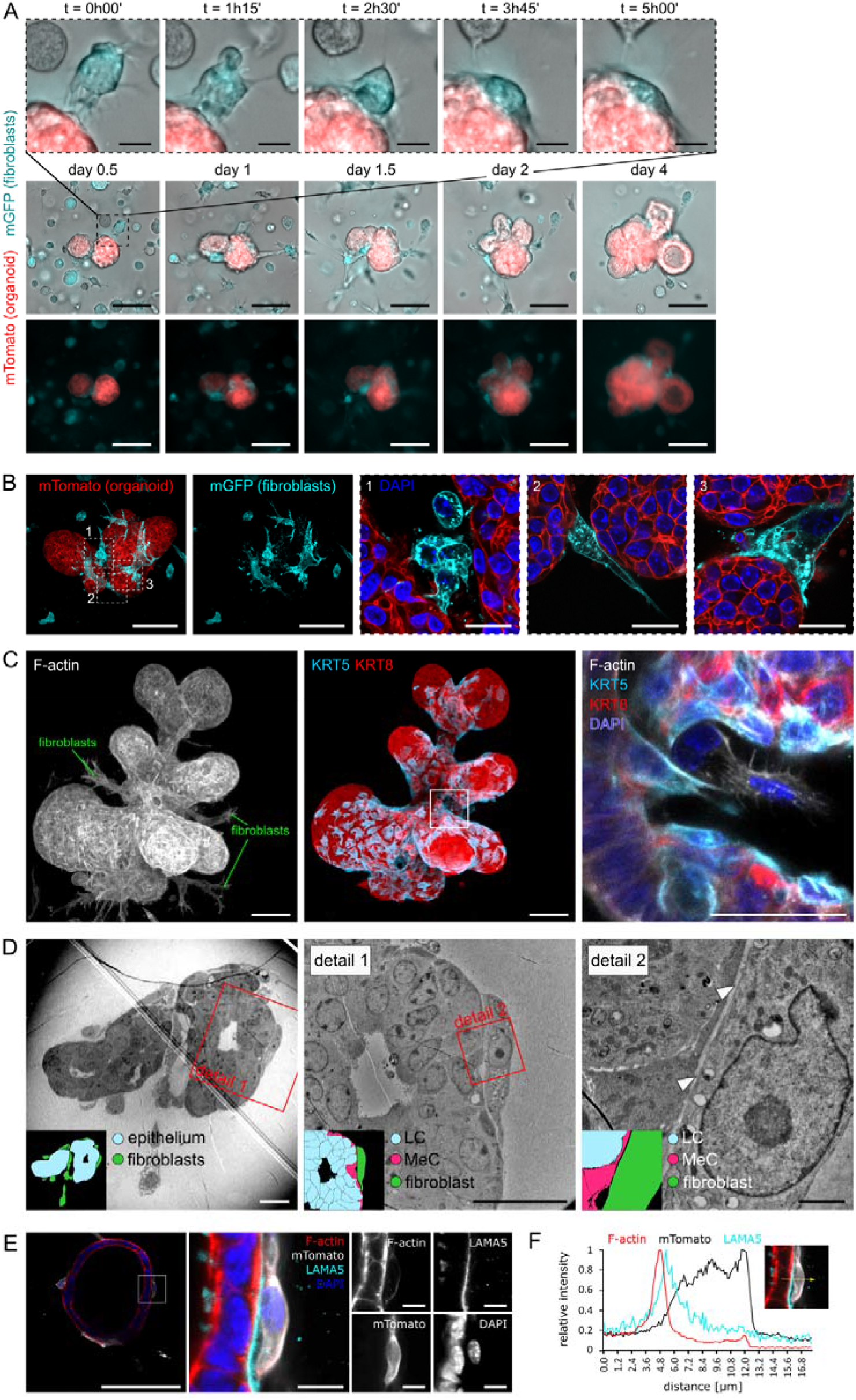
Fibroblasts in co-cultures are in physical contact with the epithelium. **A.** Snapshots from time-lapse brightfield and fluorescence imaging of organoid (tdTomato) and fibroblast (GFP) co-culture. Scale bar: 100 μm. Top line shows detail of fibroblast-organoid close interaction. Scale bar: 20 μm. **B**. Images of the contact point between organoid (tdTomato) and fibroblasts (GFP) on day 4 of co-culture. Scale bar: 100 μm, scale bar in detail: 20 μm. **C**. Images of the contact point between luminal (KRT8) and basal (KRT5) epithelial cells and fibroblasts (F-actin) on day 4 of co-culture. Scale bar: 20 μm, scale bar in detail: 10 μm. Video projection of the fibroblast-branched organoid is presented in Suppl. Video 3. **D**. Transmission electron micrographs and scheme (inset) of the contact point between luminal (LC, blue) and myoepithelial (MeC, magenta) cells and fibroblasts (green) on day 4 of co-culture. Scale bar: 20 μm, scale bar in detail: 2 μm. **E**. Optical slice of organoid-fibroblast co-culture, laminin 5 (cyan), DAPI (blue), F-actin (red), fibroblasts were isolated from *R26-mT/mG* mice (mTomato, white). Scale bar: 100 μm, scale bar in detail: 10 μm. **F**. A representative 1D relative fluorescence intensity plot. The measurement line is depicted in yellow (right).

### Fibroblast-induced epithelial branching depends on fibroblast contractility

Based on observations from the time-lapse videos of organoids branched by fibroblasts (Suppl. Videos 1 and 2) we hypothesized that fibroblasts could constrict epithelium, folding it into branches. Immunofluorescence investigation of fibroblast-branched organoids revealed that fibroblasts in contact with the organoid formed a cellular loop, encircling the branch neck (Figure 4A), and contained F-actin cables oriented mostly perpendicularly to the branch longitudinal axis (Figure 4A). Moreover, the fibroblasts expressed phosphorylated myosin light chain 2 (P-MLC2), a marker of active non-muscle myosin II (Figure 4B). Therefore, we examined the involvement of fibroblast contractility in fibroblast-induced organoid branching using small molecule inhibitors of non-muscle myosin II (blebbistatin) or ROCK1/2 (Y27632), two major nodes of cell contractility. The contractility inhibitors abrogated branching in co-cultures but did not inhibit organoid budding induced by exogenous FGF2 (Figure 4C, D; ROCK inhibition in FGF2-induced organoids even led to hyperbranched phenotype as previously described (Ewald et al., 2008)). Similarly, in the MCF7-ras spheroid co-culture model, spheroid budding was inhibited by addition of contractility inhibitors (Suppl. Figure 2A-D).

**Figure 4.**
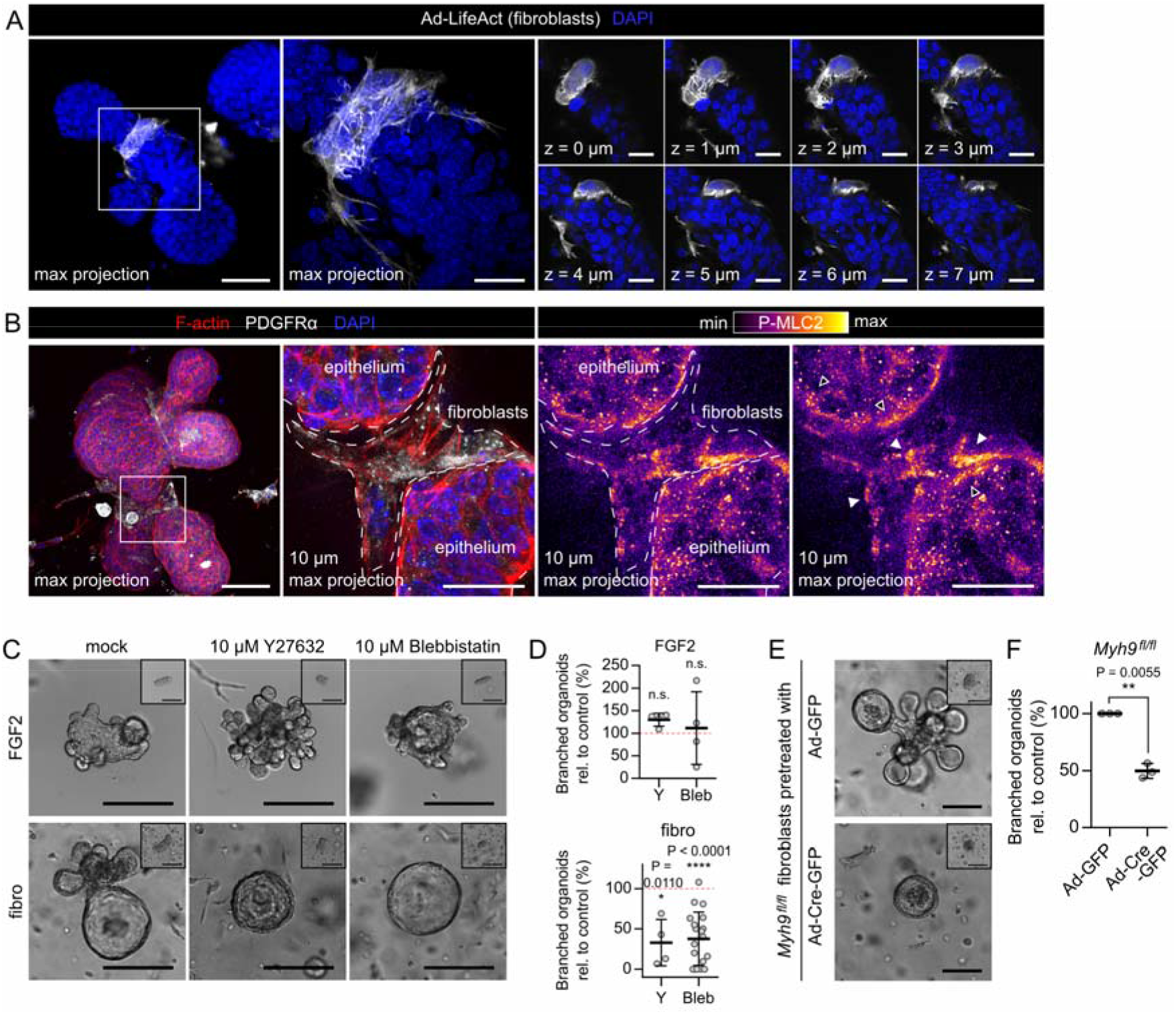
Fibroblast-induced branching requires fibroblast contractility. **A.** Maximum projection (left), detailed maximum projection (middle) and detail optical sections (right) of organoid branch induced in co-culture with fibroblasts (day 4). DAPI (blue), fibroblasts prelabeled with Ad-LifeAct-GFP (white). Scale bar: 50 μm, scale bar in detail: 20 μm. **B**. Full maximum projection (left) and 10 μm maximum projection (both middle and right) of organoid co-cultured with fibroblasts. F-actin (red), PDGFRα (white), DAPI (blue) and phosphorylated myosin light chain 2 (P-MLC2, fire LUT). Scale bar: 50 μm, scale bar in detail: 20 μm. **C**. Images of organoids on day 5 of culture (with day 0 insets) with FGF2 or with fibroblasts and treated with mock (DMSO) or contraction inhibitors (ROCK1/2 inhibitor Y27632 or non-muscle myosin II inhibitor blebbistatin). Scale bar: 100 μm. **D**. The plots show organoid branching with contraction inhibitors as mean ± SD. Statistical analysis: Multiple t-tests between each treatment and the mock-treated control; n = 4-18 (each dot represents a biologically independent experiment), N = 20 organoids per experiment. E. Images of organoids on day 5 of culture (with day 0 insets) co-cultured with control or *Myh9* knock-out fibroblasts. Scale bar: 100 μm. Videos from the 5-day experiment are presented in Suppl. Video 5. **F**. The plot shows organoid branching with mean ± SD. Statistical analysis: two-tailored paired t-test; n = 3 (each dot represents a biologically independent experiment), N = 20 organoids per experiment.

Because exogeneous treatment with pharmacological inhibitors in the culture medium affects both epithelial cells and fibroblasts, we genetically targeted exclusively in fibroblasts the contractility machinery gene myosin heavy chain 9 (*Myh9*), one of the two non-muscle myosin II heavy chain genes expressed in mammary fibroblasts (Suppl. Figure 3A, B). Both siRNA-mediated *Myh9* knockdown in wild-type fibroblasts and adenoviral Cre-mediated knock-out in *Myh9^fl/fl^* fibroblasts led to a decrease of organoid branching in co-cultures (Figure 4E, F; Suppl. Figure 4A-E; Suppl. Videos 4 and 5). These results demonstrate that fibroblast contractility is required for organoid branching in co-cultures.

### Fibroblast-induced epithelial branching requires epithelial proliferation

Mesenchymal cell tension coupled with expansion of underlying epithelium has been recently shown to cause corrugation of lung in lizard (Palmer et al., 2021). Similarly, contractile fibroblasts wrapped in loops around an expanding epithelial organoid/spheroid could be causing the branched outcome in our co-culture models. One of the common mechanisms of epithelial expansion is cell division; therefore, we characterized proliferation in the organoid-fibroblast co-culture model by both immunodetection for proliferation marker protein KI67 (Figure 5A) and by EdU incorporation (Figure 5B). EdU incorporation revealed proliferation also in fibroblast-free culture, suggesting that insulin-containing basal medium promotes cell division in organoids (Suppl. Figure 5A-C). However, we have observed regionalization of proliferation in organoids co-cultured with fibroblast with the most proliferative cells present in the neck of the branch and less at the branch tip (Figure 5C).

**Figure 5.**
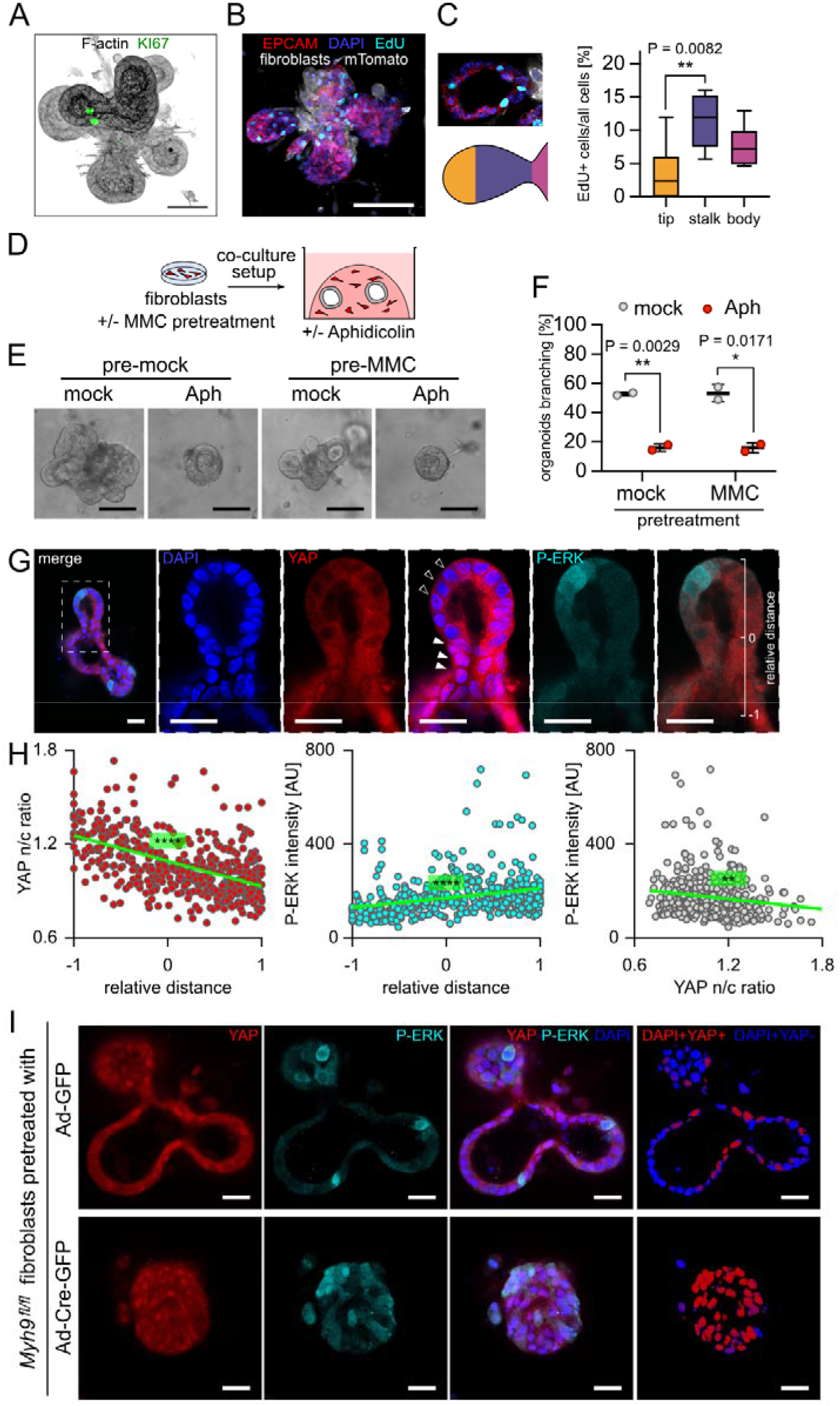
Epithelial mechanisms required for fibroblast-induced branching. **A.** F-actin (black) and KI67 (green) staining in organoids cultured with fibroblasts, day 4 of culture. Scale bar: 50 μm. The panel is related to Suppl. Figure 5A. **B**. Organoid on day 4 of co-culture, EdU administered 2 h pre-fix, EPCAM (red), DAPI (blue), EdU (cyan), fibroblasts were isolated from *R26-mT/mG* mice (mTomato, white). Scale bar: 100 μm. The panel is related to Suppl. Figure 5B. **C.** Optical section of a branch from **B** (top), a scheme of branch regions (bottom), and quantification of percentage of EdU+ cells from all cells in different branch regions. The box and whiskers plot shows minimum, median, and maximum values, and second and third quartiles of data distribution. Statistical analysis: Multiple t-tests. **D**. A scheme of the proliferation-inhibition experiment. **E**. Co-cultures at day 5, fibroblasts pretreated with +/− mitomycin C (MMC), co-cultures treated with +/− aphidicolin (Aph). Scale bar: 100 μm. **F**. Quantification of the percentage of branched organoids from experiment in **E**. The plot shows mean ± SD, each dot represent biologically independent experiment, n = 2, N = 51-77 organoids per sample, statistical analysis: t-test. **G**. Staining of YAP and phosphorylated ERK (P-ERK) in an organoid co-cultured with fibroblasts. Scale bar: 20 μm. **H.** Quantification of YAP nuclear/cytoplasmic signal ratio and P-ERK signal intensity. Relative distance −1 is branch root, +1 is branch tip. Each dot represents a single cell, n = 436 cells from 19 branches of 10 organoids. Statistical analysis: Linear regression. I. Staining of YAP and phosphorylated ERK (P-ERK) in organoids co-cultured with control or *Myh9* knock-out fibroblasts. Scale bar: 200 μm.

To test whether epithelial proliferation (and thus expansion) plays a role in organoid branching in co-cultures, we inhibited cell proliferation using aphidicolin (DNA polymerase inhibitor), upon which we observed a severe defect in organoid branching (Figure 5D-F). To test for the possibility that the observed effect could be caused by inhibition of fibroblast proliferation, we performed the experiment also with fibroblasts pretreated with mitomycin C, an irreversible DNA synthesis blocker (Figure 5D). The pretreatment of fibroblast with mitomycin C had no effect on the result (Figure 5D-F), demonstrating that fibroblast proliferation is dispensable while epithelial proliferation is necessary for organoid branching in co-cultures. In concordance with the results from organoid co-cultures, in the MCF7-ras spheroid co-culture model the blockage of spheroid proliferation by mitomycin C pretreatment of spheroids decreased spheroid size expansion and dramatically decreased branching of the MCF7-ras spheroids (Suppl. Figure 6A, B).

The regionalization of epithelial proliferation (Figure 5C) suggested an epithelial signaling response to fibroblast contact. Therefore, we examined the subcellular localization of yes associated protein (YAP), a mechano-sensor that in a resting cell resides in the cytoplasm but translocates to the nucleus upon mechanical stress (Panciera et al., 2017), and phosphorylated (active) ERK (P-ERK), the regulator of mammary epithelial branching (Ender et al., 2022; Huebner et al., 2016) as two candidates. We found that YAP specifically accumulated in the nuclei of epithelial cells in the neck region of epithelial branch (Figure 5G), indicating that the contact with contractile fibroblasts can be sensed by the underlying epithelial cells as a mechanical stress. Moreover, we found that P-ERK was localized inversely to YAP (Figure 5G, H). To test the effect of fibroblast contractility on epithelial YAP end ERK signaling, we stained YAP and P-ERK also in co-cultures with *Myh9* knockout fibroblasts. Knockout of *Myh9* in fibroblasts prevented YAP and P-ERK gradient formation (Figure 5I), indicating that contact with contractile fibroblasts regulate YAP and P-ERK epithelial distribution. Together, our data suggest that mechanical constriction by contractile fibroblasts is actively sensed by mammary epithelial cells, similarly to what was recently observed in intestinal organoids (Pérez-González et al., 2021) with the interesting difference that in our heterotypic co-culture system, the curvature of the epithelium depends on physical contact with contractile stromal cells.

## Discussion

Mechanical forces are an integral part and a driving factor of tissue morphogenesis. However, the sources of mechanical forces in different tissues are still unclear and little is understood of how force sensing is translated into cell fate during organ formation. Our work reveals the critical role of fibroblast-derived mechanical forces in regulation of mammary epithelial branching morphogenesis. It demonstrates that mammary fibroblasts generate mechanical forces via their actomyosin apparatus and transmit them to the epithelium, which leads to epithelial deformation and patterning of epithelial intracellular signaling, resulting in epithelial folding into branched structures.

### Fibroblast-generated mechanical forces as part of complex tissue mechanics

The role of intraepithelial forces in morphogenetic processes involving tissue folding, such as gastrulation, tubulogenesis, or buckling has been long recognized and intensively studied (Heisenberg and Bellaïche, 2013). Similarly, the instructive role of mechanical properties and 3D organization of the ECM in determination of cell fate and behavior during organ formation, including mammary epithelial branching morphogenesis, has been well established (Bonnans et al., 2014; Brownfield et al., 2013; Nguyen-Ngoc and Ewald, 2013). However, the evidence for regulation of epithelial morphogenesis by mechanical stimuli from mesenchymal cells was discovered only recently and has been scarce, limited to the morphogenesis of feather buds in chick skin by mechanically active dermal cells (Shyer et al., 2017), gut villification (Shyer et al., 2013) and lung epithelial bifurcation and alveologenesis induced by smooth muscle cells or myofibroblasts (Goodwin et al., 2019; Kim et al., 2015; Li et al., 2020a; Palmer et al., 2021).

### The mechanism of fibroblast-induced mammary morphogenesis: connection to ECM remodeling

It was previously proposed that mechanical forces generated by mesenchymal/stromal cells regulate epithelial morphogenesis indirectly via changes of ECM mechanics, including collagen I remodeling in embryonic gut (Hughes et al., 2018) and postnatal mammary gland (Koledova et al., 2016; Peuhu et al., 2017; Sumbal et al., 2020), or elastin deposition in lung (Li et al., 2018). However, while not excluding contribution of such mechanism to mammary epithelial branching *in vivo*, our investigations *in vitro* in organoid-fibroblast co-cultures devoid of collagen I demonstrate that collagen I fibers are not required for induction of epithelial folding by fibroblast contractility. Mammary fibroblasts form direct, highly dynamic contacts with mammary epithelial cells and induce a mechanosensitive response in the epithelium, resulting in patterning of key morphogenetic regulators YAP and ERK. The direct contact between mammary fibroblasts and epithelial cells *in vivo* could be enabled by immature, thin basement membrane of terminal end buds (Silberstein and Daniel, 1982), the highly proliferative epithelial structures, which drive postnatal mammary branching morphogenesis, and active remodeling of ECM by matrix metalloproteinases produced by both epithelial cells and fibroblasts (Feinberg et al., 2018), which is essential for mammary branching morphogenesis (Fata et al., 2004; Feinberg et al., 2018). Our work does not exclude the importance of ECM remodeling by fibroblast mechanical forces in epithelial branching. We speculate that *in vivo* the highly dynamic mechanically active fibroblasts could initiate formation of epithelial clefts and further reinforce them by subsequent deposition and remodeling of ECM.

### The mechanism of fibroblast-induced mammary morphogenesis: requirement of paracrine signaling

Importantly, our results do not rule out the importance of fibroblast-secreted factors in mammary epithelial morphogenesis. However, we demonstrate that paracrine signals are not sufficient to drive organoid branching in the 3D *in vitro* cultures of organoids with fibroblasts without addition of any branching-inducing growth factors and show the importance of fibroblast-epithelium contact, so short-distance paracrine or juxtacrine signals could be important in the process. Several growth factors, including FGF2, FGF7, EGF or TGFα can induce organoid branching in the absence of fibroblasts when added to the medium in nanomolar concentrations (Sternlicht et al., 2005). However, the evidence for requirement of those growth factors’ expression in mammary fibroblasts for mammary epithelial branching *in vivo* is missing.

### The mechanism of fibroblast-induced mammary morphogenesis: epithelial response

Our work reveals that mechanical strain imposed on mammary epithelial cells by fibroblasts results in epithelial folding with negative curvature in the epithelial-fibroblast contact points, which show low ERK activity, and positive curvature in the branch tips, which show high ERK activity. This distribution of active ERK is in line with the role of ERK in mammary branch elongation (Huebner et al., 2016) and with a recent study indicating that ERK is specifically activated in epithelia with a positive curvature in the developing lung (Hirashima and Matsuda, 2021). While the mechanisms of epithelial folding, ERK spatial pattern formation, and their interplay during fibroblast-induced mammary epithelial morphogenesis remain to be investigated, they could include ERK-mediated mechanochemical waves as described during 2D collective cell migration (Hino et al., 2020).

Importantly, the direct interactions between mammary epithelium (including both organoids from normal epithelium and spheroids from breast cancer cells) and fibroblasts do not lead to invasive dissemination of epithelial cells, unlike in co-cultures of squamous cell carcinoma with cancer-associated fibroblasts (CAFs) (Labernadie et al., 2017). Interestingly, a recent study described mechanical compression of intestinal tumors by CAFs forming a mechanically active tumor capsule (Barbazan et al., 2021), providing further evidence for context-dependent employment of fibroblast-derived mechanical forces in tissue morphogenesis and tumorigenesis.

### Fibroblasts as central regulators of epithelial morphogenesis and homeostasis: evidence for mechanically active fibroblasts in vivo

Fibroblasts accompany mammary epithelial cells from early development through homeostasis to aging and disease and employ different functions to meet epithelial needs (Sumbal et al., 2020). The multiple fibroblast functions are facilitated by fibroblast heterogeneity, which has only recently begun to be resolved using single-cell RNA sequencing approaches (Li et al., 2020b; Yoshitake et al., 2022). These studies confirmed well-established fibroblast roles in epithelial development and tissue homeostasis via production of paracrine signals and ECM, and fibroblast roles in regulation of immune landscape of the mammary gland. Though they did not detect mechanically active fibroblasts. However, these studies included only adult and aged mammary glands and omitted puberty, the stage of active mammary epithelial branching morphogenesis, thus the *in vivo* existence of mechanically active mammary fibroblasts in normal mammary gland requires further investigation. The presence and function of mechanically active fibroblasts (contractile fibroblasts expressing αSMA, myofibroblasts) has been well documented in other developing organs (Li et al., 2018) and in multiple tumor types, including mammary carcinomas (Costa et al., 2018; Sahai et al., 2020).

In conclusion, we find that fibroblasts drive branching morphogenesis of the mammary gland by exerting mechanical forces on epithelial cells. It is conceivable that such conserved mechanism could be used to regulate morphogenesis of other branched organs, providing a comprehensive understanding of overlapping but divergent employment of mechanically active fibroblasts in developmental versus tumorigenic programs.

## Materials and Methods

### Animals

All procedures involving animals were performed under the approval of the Ministry of Agriculture of the Czech Republic (license # MSMT-9232/2020-2), supervised by the Expert Committee for Laboratory Animal Welfare of the Faculty of Medicine, Masaryk University, at the Laboratory Animal Breeding and Experimental Facility of the Faculty of Medicine, Masaryk University (facility license #58013/2017-MZE-17214), or under the approval of the ethics committee of the Institut Curie and the French Ministry of Research (reference #34364-202112151422480) in the Animal Facility of Institut Curie (facility license #C75–05–18). ICR mice were obtained from the Laboratory Animal Breeding and Experimental Facility of the Faculty of Medicine, Masaryk University. *R26-mT/mG* (Muzumdar et al., 2007) and *Acta2-CreERT2* mice (Wendling et al., 2009) were acquired from the Jackson Laboratories. LifeAct-GFP mice (Riedl et al., 2010) were created by Wedlich-Söldner team, *Myh9*^fl/fl^ mice (Conti et al., 2004) were kindly provided by Dr. Sara Wickström. Transgenic animals were maintained on a C57/BL6 background. Experimental animals were obtained by breeding of the parental strains, the genotypes were determined by genotyping. The mice were housed in individually ventilated or open cages, all with ambient temperature of 22°C, a 12□h:12□h light:dark cycle, and food and water *ad libitum*. Female 6-8 weeks old virgin mice were used in the experiments. Mice were euthanized by cervical dislocation and mammary gland tissues were collected immediately.

### Primary mammary organoid and fibroblast isolation and culture

Primary mammary fibroblasts and organoids were isolated from 6-8 weeks old female virgin mice (ICR, unless otherwise specified) as previously described (Koledova, 2017). The mammary glands were chopped and partially digested in a solution of collagenase and trypsin [2 mg/ml collagenase A, 2 mg/ml trypsin, 5 μg/ml insulin, 50 μg/ml gentamicin (all Merck), 5% fetal bovine serum (FBS; Hyclone/GE Healthcare) in DMEM/F12 (Thermo Fisher Scientific)] for 30 min at 37°C. Resulting tissue suspension was treated with DNase I (20 U/ml; Merck) and submitted to five rounds of differential centrifugation (450 × g for 10 s) to separate epithelial (organoid) and stromal fractions. The organoids were resuspended in basal organoid medium [1× ITS (10 μg/ml insulin, 5.5 μg/ml transferrin, 6.7 ng/ml sodium selenite), 100 U/ml of penicillin, and 100 μg/ml of streptomycin in DMEM/F12] and kept on ice until co-culture setup. The cells the from stromal fraction were pelleted by centrifugation, suspended in fibroblast cultivation medium (10% FBS, 1× ITS, 100 U/ml of penicillin, and 100 μg/ml of streptomycin in DMEM) and incubated on cell culture dishes at 37°C, 5% CO2 for 30 min. Afterwards the unattached (non-fibroblast) cells were washed away, the cell culture dishes were washed with PBS and fresh fibroblast medium was provided for the cells. The cells were cultured until about 80% confluence. During the first cell subculture by trypsinization, a second round of selection by differential attachment was performed, when the cells were allowed to attach only for 15 min at 37°C and 5% CO_2_. The fibroblasts were expanded and used for the experiments until passage 5.

To inhibit fibroblast proliferation for specific assays, the fibroblasts were treated with 10 μg/ml mitomycin C in fibroblast medium for 3 h at 37°C, 5% CO_2_. Afterwards the fibroblasts were washed three times with PBS and one time with basal organoid medium, trypsinized and used to set up co-cultures.

To prepare fibroblast-conditioned medium, the fibroblasts were seeded in cell culture dishes in fibroblast medium and the next day, the cells were washed three times with PBS and incubated with basal organoid medium for 24 h. Afterwards the medium was collected from the dishes, sterile-filtered through a 0.22 μm filter, and used immediately in the experiment, or aliquoted, stored at −20°C and used within 5 days of conditioned medium preparation.

### 3D culture of mammary organoids and fibroblasts

3D culture of mammary organoids and fibroblasts was performed as previously described (Koledova and Lu, 2017). Freshly isolated mammary organoids were embedded in Matrigel either alone (300 organoids in 45 μl of Matrigel per well) or with 5×10^4^ mammary fibroblasts per well and plated in domes in 24-well plates. For transwell experiments, organoids were plated in domes in the transwell (8 μm pore size, Falcon-Corning), fibroblasts were plated in lower chamber. After setting the gel for 45-60 min at 37°C, the cultures were overlaid with basal organoid medium (1× ITS, 100 U/ml of penicillin, and 100 μg/ml of streptomycin in DMEM/F12), not supplemented or supplemented with growth factors [2.5 nM FGF2 (Enantis)] or small molecule inhibitors (Suppl. Table 1) according to the experiment. The cultures were incubated in humidified atmosphere of 5% CO_2_ at 37°C on Olympus 1X81 microscope equipped with Hamamatsu camera and CellR system for time-lapse imaging. The organoids/co-cultures were photographed every 60 min for 5 days with manual refocusing every day (high-detail imaging) or photographed only once per day for 5 days (low-detail imaging). The images were exported and analyzed using Image J. Organoid branching was evaluated from videos and it was defined as formation of a new bud/branch from the organoid. Organoids that fused with another organoid or collapsed after attachment to the bottom of the dish were excluded from the quantification.

For fluorescent time-lapse imaging, organoids were isolated from *R26-mT/mG* mammary glands on day of the experiment. Fibroblasts were isolated from *Acta2-CreERT2;mT/mG* mice, cultured to passage 2-3 and induced *in vitro* by 0.5 mM 4-OH-tamoxifen (Sigma) treatment for 4 days prior trypsinization and experimental use. Before experiment, the GFP fluorescence of fibroblasts was assessed using a microscope and when it was > 95%, the cells were used for co-culture. Co-cultures were seeded on coverslip-bottom 24-well plate (IBIDI) and imaged on Cell Discoverer 7 equipped with PLAN-APOCHROMAT 20x/0.95 autocorr with 0.5× magnification lens. GFP was imaged with 470/40 excitation, 525/50 emission, tdTomato was imaged with 545/25 excitation, 605/70 emission filter (all Zeiss). The samples were incubated in a humidified atmosphere of 5% CO_2_ at 37°C during the imaging.

### 3D culture of spheroids and fibroblasts

MCF7-ras cells (kindly provided by Dr. Ula Polanska) were expanded in DMEM/F12 supplemented with 10% FBS, 100 U/ml of penicillin, and 100 μg/ml of streptomycin and incubated in non-adherent PolyHEMA-coated dish overnight to form spheroids. Next day, the spheroids were embedded either alone (200 spheroids in 45 μl of Matrigel per well) or with 5×10^4^ mammary fibroblasts per well and plated in domes in 24-well plates. After setting the gel for 45-60 min at 37°C, the cultures were overlaid with basal organoid medium, supplemented with growth factors [2.5 nM FGF2 (Enantis) or EGF (Peprotech)] small molecule inhibitors (Suppl. Table 1) according to the experiment. The (co-)cultures were incubated in a humidified atmosphere of 5% CO_2_ at 37°C on Olympus IX81 microscope equipped with Hamamatsu camera and CellR system for time-lapse imaging and photographed every 60 min for 5 days with manual refocusing every day (high-detail imaging) or photographed only once per day for 5 days (low-detail imaging). The images were exported and analyzed using Image J. Spheroid budding was evaluated from the videos and it was defined as formation of a new bud from the spheroid. Spheroids that fused with other spheroids were excluded from the quantification.

### Knockdown and knockout of *Myh9* in mammary fibroblasts

For *Myh9* knockdown, the pre-designed Silencer Select siRNAs against *Myh9* (IDs s70267 and s70268, Myh9si#1 and Myh9si#2, respectively) and the scrambled negative control siRNA (Silencer Select negative control or Stealth negative control siRNA), all from Thermo Fisher Scientific, were transfected into wild type (ICR) fibroblasts with Lipofectamine 3000 Reagent (Thermo Fisher Scientific) according to manufacturer’s instructions at 20 nM siRNA. For *Myh9* knockout, *Myh9^fl/fl^* fibroblasts were transduced with adenoviruses Adeno-Cre-GFP (Ad-Cre-GFP) or Adeno-GFP (Ad-GFP) from Vector Biolabs at 200 MOI for 4 h. Next day, the transfected/transduced fibroblasts were put in co-culture with organoids and submitted to time-lapse imaging. A part of the fibroblasts was further cultured and knockdown/knockout efficiency was determined 72 h after transfection/transduction by qPCR analysis of *Myh9* mRNA levels, normalized to housekeeping genes *Actb* and *Eef1g*, and by immunostaining for MYH9.

### LifeAct adenoviral transduction

For imaging experiments with LifeAct, fibroblasts were infected with LifeAct adenoviral particles (IBIDI) according to the manufacturer’s instructions prior to co-culture set-up. Briefly, the adenovirus particles were reconstituted in fibroblast cultivation medium at concentration of 500 MOI and incubated with adherent fibroblasts at 37°C for 4 h. After that, adenovirus-containing medium was washed out, and the cells were kept overnight in fibroblast cultivation medium. The next day, GFP fluorescence was checked under the microscope and when > 50% of cells appeared green, fibroblasts were used for co-culture.

### Immunofluorescence staining of 2D fibroblasts

For immunofluorescent analysis, fibroblasts were cultured directly on glass coverslips, fixed with 10% neutral buffered formalin, permeabilized with 0.05% Triton X-100 in PBS and blocked with PBS with 10% FBS. Then the coverslips were incubated with primary antibodies (Suppl. Table 2) for 2 h at RT or overnight at 4°C. After washing, the coverslips were incubated with secondary antibodies and phalloidin AlexaFluor 488 (Suppl. Table 2) for 2 h at RT. Then the coverslips were washed, stained with DAPI (1 μg/ml; Merck) for 10 min and mounted in Mowiol (Merck). The cells were photographed using Axio Observer Z1 microscope with laser scanning confocal unit LSM 800 with 405, 488, 561 and 640 nm lasers, GaAsp PMT detector and objective Plan-Apochromat 40x /1.20 and C-Apochromat 63x /1.20 with water immersion (all Zeiss). The brightness of each channel was linearly enhanced in Zen Blue software (Zeiss) and pictures were cropped to final size in Photo Studio 18 (Zoner).

### Immunofluorescence staining of 3D co-cultures

For immunofluorescent analysis of 3D co-cultures, the co-cultures were fixed with 10% neutral buffered formalin, washed, and stored in PBS. Next, organoid co-cultures were stained according to the droplet-based method as described (Sumbal and Koledova, 2022). Briefly, the fixed co-cultures were placed on stereoscope (Leica FM165C) and pieces containing an organoid with approximately 100 μm of surrounding Matrigel with fibroblasts were manually cut out with 25G needles and moved on parafilm-covered cell culture dish for staining. All the staining steps were done on the parafilm in 20 μl drops and all solutions were changed under direct visual control using the stereoscope. The co-cultures were permeabilized with 0.5% Triton X-100 in PBS, blocked with 8% FBS and 0.1% Triton X100 in PBS (3D staining buffer, 3SB) and incubated with primary antibodies (Suppl. Table 2) in 3SB over 1-3 nights at 4°C. Then the co-cultures were washed for 3h with 0.05% Tween-20 in PBS and incubated with secondary antibodies, phalloidin AlexaFluor 488 (Suppl. Table 2) and DAPI (1 μg/ml; Merck) in 3SB over 1-2 nights at 4°C in dark. Then the co-cultures were washed for 3h with 0.05% Tween-20 in PBS, cleared with 60% glycerol and 2.5 M fructose solution overnight at RT in dark and mounted between slide and coverslip with double-sided tape as a spacer. The co-cultures were imaged using inverted microscope Axio Observer 7 with laser scanning confocal unit LSM 880 with 405, 488, 561 and 633 nm lasers, GaAsp PMT spectral detector and objective C-Apochromat 40x /1.20 or C-Apochromat 63x / 1.20 with water immersion (all Zeiss). The co-cultures were photographed either as one optical slice or like 3D z-stacks of various z-step as required per experiment. The brightness of each channel was linearly enhanced in Zen Blue software (Zeiss) and pictures were cropped to final size in Photo Studio 18 (Zoner). Branch morphometry was manually analyzed from confocal images using ImageJ measure length function, measuring the thickness perpendicular to the long axis of the branch, in the central z-plane.

### Image analysis of signal distribution

The analysis of P-ERK intensity and YAP nuclear to cytoplasmic ratio was done in ImageJ (NIH). Cells in optical section in the middle of an organoid branch were manually annotated and segmented for target protein signal (YAP and P-ERK channels) and nuclei (DAPI channel) and density of pixels in YAP and P-ERK channels in the regions of interest (ROIs) was measured. The nuclear to cytoplasmic ratio of YAP was calculated in excel (Microsoft). The spatial information of each ROI was manually measured on a line parallel to the branch longitudinal axis and normalized, with the value “1” set for the tip of the branch and the value “-1” set for the root of the branch. The graphs and linear regression line were created in Prism 6 (GraphPad). Colocalization analysis of YAP and DAPI channels was done in Zen Black (Zeiss) and presented as color-coded (blue DAPI+YAP- and red DAPI+YAP+). The same cut-off for the colocalization analysis was applied for all images from the same experiment.

### Real-time quantitative PCR (qPCR)

RNA from fibroblasts was isolated using RNeasy Mini Kit (Qiagen) according to the manufacturer’s instruction. RNA concentration was measured using NanoDrop 2000 (Thermo Fisher Scientific). RNA was transcribed into cDNA by using Transcriptor First Strand cDNA Synthesis Kit (Roche) or TaqMan Reverse Transcription kit (Life Technologies). Real-time qPCR was performed using 5 ng cDNA, 5 pmol of the forward and reverse gene-specific primers each (primer sequences are shown in Suppl. Table 3) in Light Cycler SYBR Green I Master mix (Roche) on LightCycler 480 II (Roche). Relative gene expression was calculated using the ΔΔCt method and normalization to two housekeeping genes, β- actin (*Actb*) and eukaryotic elongation factor 1 γ (*Eef1g*).

### Transmission electron microscopy

The 3D co-cultures were fixed with 3% glutaraldehyde in 100 mM sodium cacodylate buffer, pH 7.4 for 45 min, postfixed in 1% OsO_4_ for 50 min, and washed with cacodylate buffer. After embedding in 1% agar blocks, the samples were dehydrated in increasing ethanol series (50, 70, 96, and 100%), treated with 100% acetone, and embedded in Durcupan resin (Merck). Ultrathin sections were prepared using LKB 8802A Ultramicrotome, stained with uranyl acetate and Reynold’s lead citrate (Merck), and examined with FEI Morgagni 286(D) transmission electron microscope. The cells in the schematics were segmented manually.

### Statistical analysis

Sample size was not determined *a priori* and investigators were not blinded to experimental conditions. Statistical analysis was performed using GraphPad Prism software. Student’s t-test (unpaired, two-tailed) was used for comparison of two groups. Bar plots were generated by GraphPad Prism and show mean ± standard deviation (SD). *P < 0.05, **P < 0.01, ***P < 0.001, ****p < 0.0001. The number of independent biological replicates is indicated as n.

## Acknowledgements

We are grateful to Danijela Matic Vignjevic for critical review of the manuscript, to Denisa Belisova for mouse husbandry, and to Maria Luisa Martin Faraldo for the LAMA5 antibody. We acknowledge the core facility CELLIM of CEITEC, supported by the Czech-Biolmaging large Rl project (LM2018129 funded by MEYS CR), for their support with obtaining scientific data presented in this paper. We gratefully acknowledge the Cell and Tissue Imaging Platform (PICT-IBiSA) at Institut Curie, member of the French National Research Infrastructure France-BioImaging (ANR-10-INBS-04). We are thankful to Enantis for providing FGF2-wt. J.S. is supported by Barrande Fellowship (Ministry of Education, Youth and Sports) and FRM FDM202106013570, and by Brno PhD. Talent Scholarship, funded by the Brno City Municipality. This work was supported by grants from the Grant Agency of Masaryk University (MU) (projects no. MUNI/G/1446/2018, MUNI/G/1775/2020 to Z.S.K., MUNI/A/1398/2021 and MUNI/A/1301/2022) and from Internal Grant Agency of Faculty of Medicine MU (MUNI/11/SUP/06/2022 to Z.S.K. and MUNI/IGA/1314/2021 to J.S.).

## Author Contributions

Z.S.K. conceptualized the study. J.S. and Z.S.K. designed the experiments. J.S. and Z.S.K. performed all experiments, data analysis, quantification, and statistical analysis. S.F. supervised and funded the experiments performed at Institut Curie. Z.S.K. and J.S. wrote the manuscript. All authors reviewed and edited the manuscript and agreed on its final version.

## Conflict of Interest

The authors declare no competing financial interests.

## Data availability

The authors declare that all data supporting the findings of this study are available within the article and its supplemental information files or from the corresponding author upon reasonable request.

## Limitations of the study

The critical experiment that demonstrates the need of fibroblasts’ physical contact with the epithelium for epithelial branching does not allow to distinguish between direct physical contact and potential juxtacrine or very short-distance paracrine signaling between the epithelium and fibroblasts, which may contribute to epithelial morphogenesis.

## Notes

### Competing Interest Statement

The authors have declared no competing interest.

### Summary of Updates

In the revised version of the manuscript, supplementary files have been added.

