## Supplemental Information for "Fibroblast-induced mammary epithelial branching depends on fibroblast contractility"

### Supplementary Tables

**Supplementary Table 1. The list of pharmacological and viral compounds.**

| Compound | Supplier | Cat. Number | Concentration used |
|---|---|---|---|
| Aphidicolin | Merck | A4487 | 1.5 µM |
| Blebbistatin | Merck | B0560 | 10 µM |
| Mitomycin C | Merck | M4287 | 10 µg/ml |
| SU5402 | Merck | SML0443 | 1 µM, 5 µM, 10 µM |
| U0126 | Bio-techne | 1144 | 0.5 µM, 1 µM, 2 µM |
| Y27632 | Merck | SCM075; Y0503 | 10 µM |
| 4-OH tamoxifen | Sigma | H7904 | 0.5 mM |
| Adeno-GFP | Vector Biolabs | 1060 | 200 MOI |
| Adeno-Cre-GFP | Vector Biolabs | 1700 | 200 MOI |
| rAVCMV-LifeAct-TagGFP2 | IBIDI | 60120 | 500 MOI |

**Supplementary Table 2. The list of detection agents used in this study.**

| Antibody | Host, class | Supplier | Cat. Number | Dilution |
|---|---|---|---|---|
| **ICC-IF** | | | | |
| MYH9 | Rabbit, polyclonal | Biolegend | 909801 | 1/1000 |
| MYH10 | Rabbit, polyclonal | Biolegend | 909901 | 1/1000 |
| *Secondary antibodies* | | | | |
| AlexaFluor 488 conjugated | Goat, polyclonal | Thermo Fisher Scientific | A-11001<br>A-11008 | 1/1000<br>1/1000 |
| Alexa Fluor 546 conjugated | Goat, polyclonal | Thermo Fisher Scientific | A-21133 | 1/1000 |
| Alexa Fluor 568 conjugated | Goat, polyclonal | Thermo Fisher Scientific | A-11011 | 1/1000 |
| Alexa Fluor 647 conjugated | Goat, polyclonal | Thermo Fisher Scientific | A-21235<br>A-21244 | 1/1000<br>1/1000 |
| **Organoid 3D staining** | | | | |
| Keratin 5 | Rabbit, polyclonal | Biolegend | 905504 | 1/250 |
| Keratin 8 | Rabbit, polyclonal | Biolegend | 904804 | 1/250 |
| KI67 | Rabbit, polyclonal | Zytomed | RBK027 | 1/300 |
| P-MYL9 (S19) | Mouse, monoclonal | Thermo Fisher Scientific | MA515163 | 1/100 |
| PDGFRα | Rabbit, monoclonal | Cell Signaling Technology | #3174 | 1/100 |
| P-ERK1/2 (T202/Y204) | Rabbit, monoclonal | Cell Signaling Technology | #4370 | 1/250 |
| YAP | Mouse, monoclonal | Santa Cruz Biotechnology | sc-101199 | 1/100 |
| Laminin α5 | Rabbit, polyclonal | (Rousselle and Aumailley, 1994) | N/A | 1/100 |
| *Secondary antibodies*<br>as above | | | | 1/800 |
| Phalloidin-AlexaFluor488 | N/A | Thermo Fisher Scientific | A12379 | 1/200 |

**Supplementary Table 3. The list of primers used for qPCR in this study.**

| Gene name | Forward primer (5'-3') | Reverse primer (5'-3') | length [bp] |
|---|---|---|---|
| ***Actb*** | GGCTGTATTCCCCTCCATCG | CCAGTTGGTAACAATGCCATGT | 154 |
| ***Eef1g*** | TTCCTGCCGGCAAGGTTCCA | TGCCGCCTCTGGCGTACTTC | 119 |
| ***Myh9*** | GGCCCTGCTAGATGAGGAGT | CTTGGGCTTCTGGAACTTGG | 106 |
| ***Myh10*** | GGAATCCTTTGGAAATGCGAAGA | GCCCCAACAATATAGCCAGTTAC | 102 |
| ***Myh14*** | CAGTGACCATGTCCGTGTCTG | CGTAGAGGAACGATTGGGCTG | 81 |

### Supplementary Figures

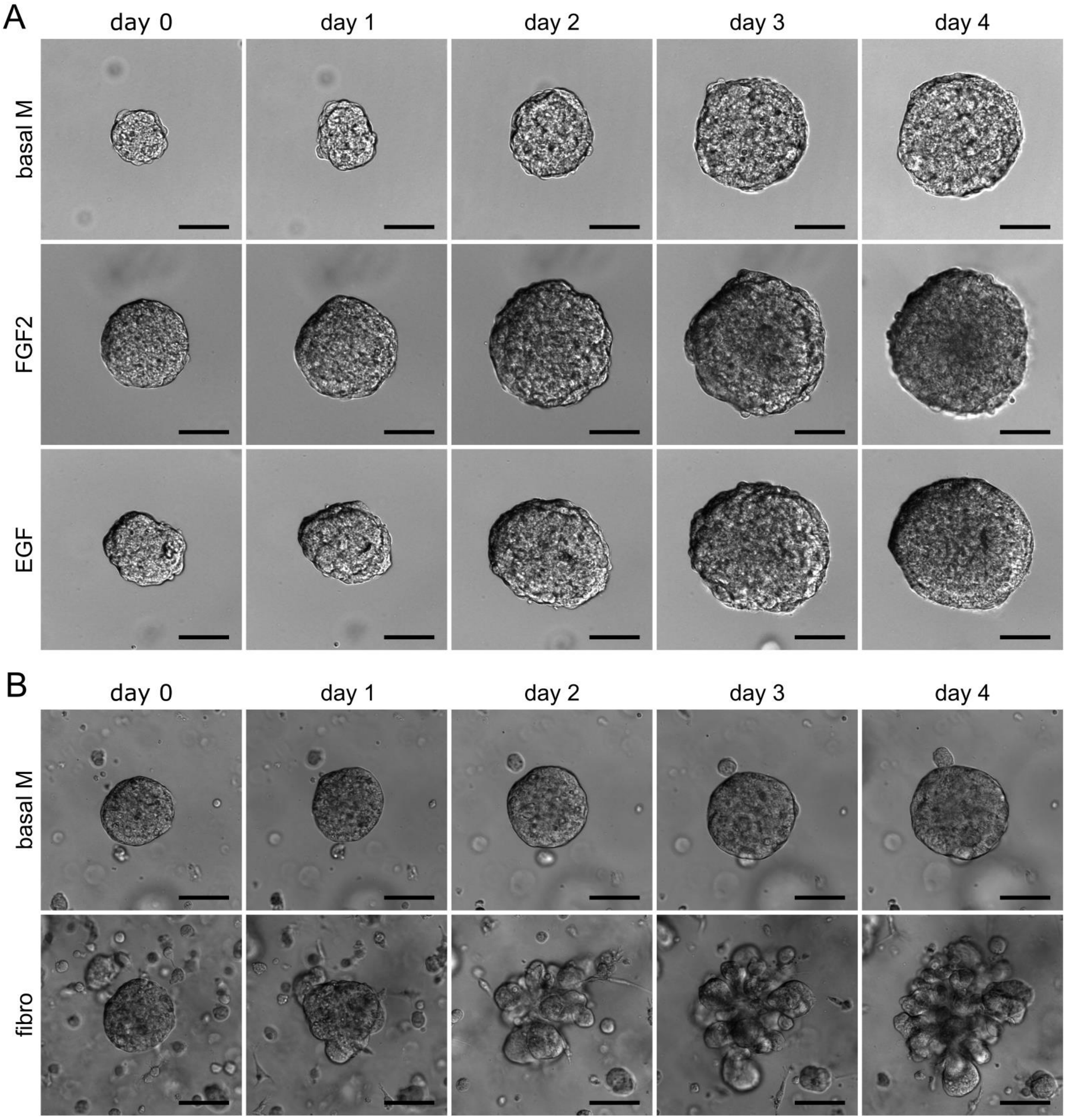


**Supplementary Figure 1. MCF7-ras spheroids do not respond to exogenous growth factors by branching**. **A.** Time-lapse snapshots of MCF7-ras spheroids cultured in basal medium with no exogenous growth factors (basal M) or with FGF2 or EGF. Scale bar: 100 μm. **B.** Time-lapse snapshots of MCF7-ras spheroids co-cultured with no stromal cells (basal M) or with fibroblasts (fibro). Scale bar: 100 μm.

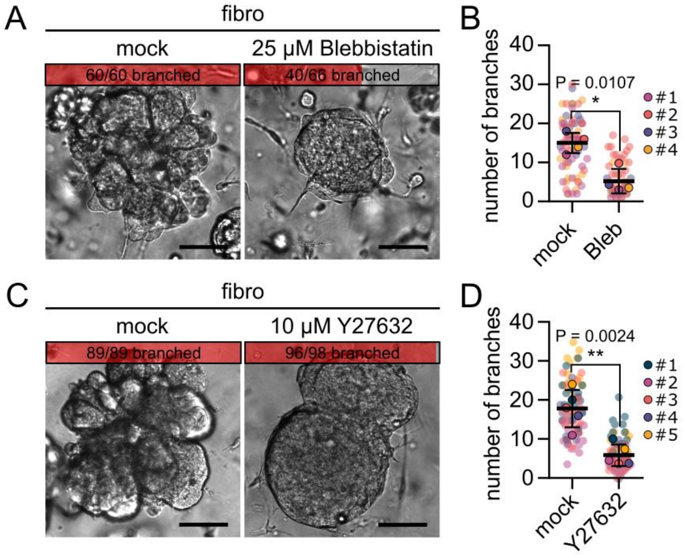


**Supplementary Figure 2. MCF7-ras spheroid budding in co-cultures requires cell contractility. A, C.** Photographs of spheroids on day 4 of co-culture with fibroblasts upon treatment with no inhibitor (mock), with Blebbistatin (Bleb, **A**) or with Y27632 (**C**). Top grey and red bars indicate proportion of branched spheroids out of all spheroids per condition. Scale bar: 100 μm. **B, D.** Quantification of number of branches/buds per branched spheroid in conditions from A. The plot shows mean ± SD, each lined dot shows mean from each experiment, each faint dot shows single spheroid measurement, n = 4 (**B**) or 5 (**D**) biologically independent experiments, N = 20 spheroids per experiment. Statistical analysis: two-tailored *t*-test.

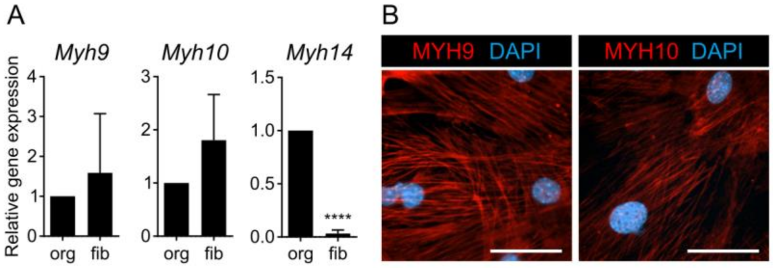


**Supplementary Figure 3. Mammary fibroblasts express MYH9 and MYH10. A.** Real-time qPCR analysis of myosin II heavy chain genes *Myh9, Myh10* and *Myh14* in mammary fibroblasts (fib) and epithelium (organoids, org). Plots show mean ± SD. Statistical analysis: two-tailored *t*-test; n = 3 independent biological samples. **B.** Representative images of MYH9 and MYH10 immunostaining in mammary fibroblasts in first passage. Scale bar: 50 μm.

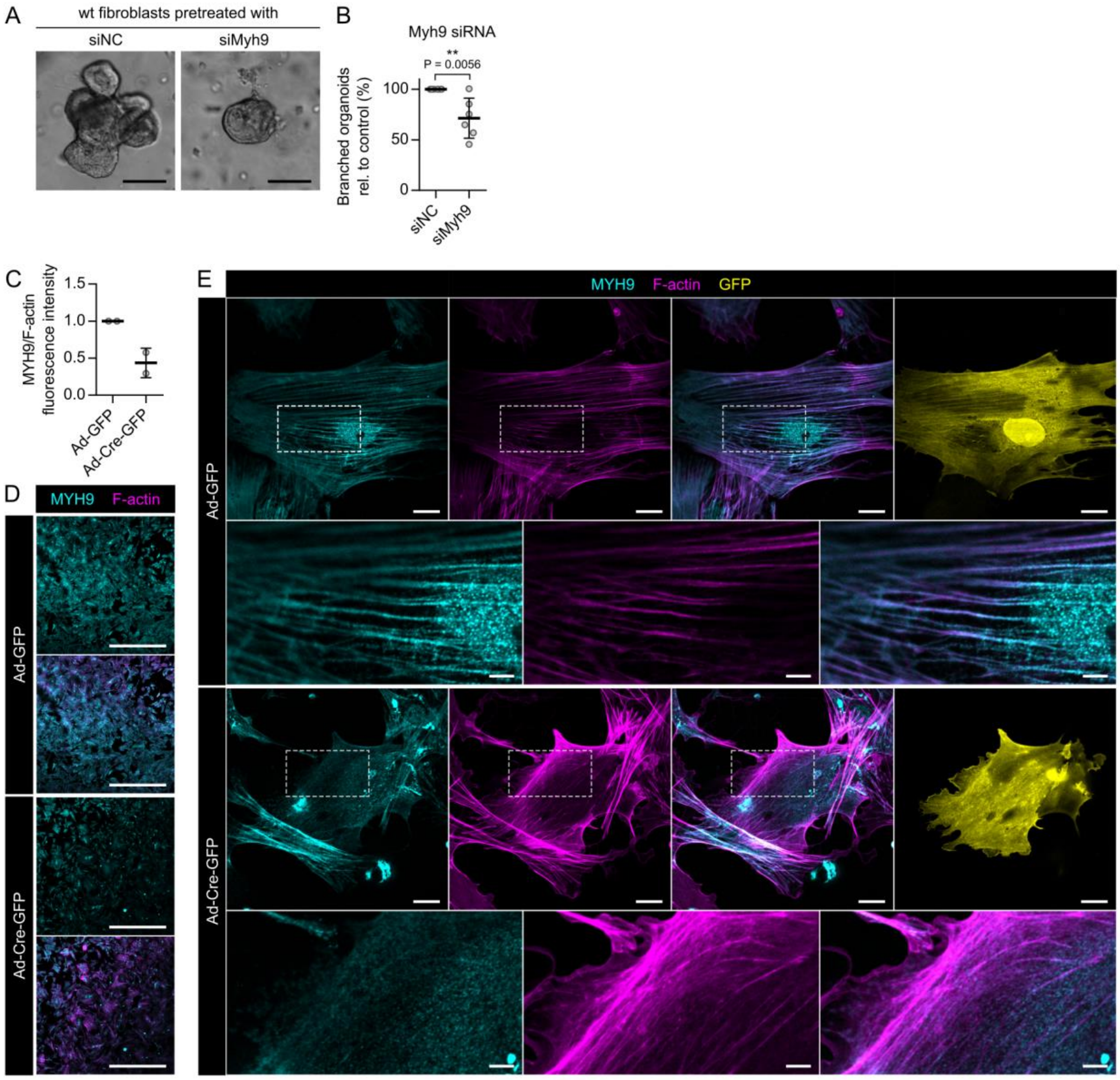

**Supplementary Figure 4. Knockdown of *Myh9* in mammary fibroblasts abrogates fibroblast-induced branching of mammary organoids. A.** Representative images (day 5 of culture) and quantification of organoid branching in co-cultures with fibroblasts pre-treated with nonsense (siNC) or *Myh9* targeting (siMyh9) siRNA. Plot indicates mean ± SD. Statistical analysis: two-tailored paired *t*-test; n = 6 independent *Myh9* knockdown experiments; N = 20 organoids per each treatment of each independent experiment. Videos from the 5-day experiment are presented in Suppl. Video 4. **B-D.** Quantification of MYH9 downregulation in *Myh9* KO fibroblasts by immunofluorescence. The plot (**B**) shows mean ± SD, n = 2 independent experiments. Representative images (**C**) show MYH9 (cyan) and F-actin (phalloidin, magenta) staining in cultured primary mammary fibroblasts from *Myh9*$^{fl/fl}$ mice, treated with adeno-GFP (Ad-GFP) or adeno-Cre-GFP (Ad-Cre-GFP) vector, including details (**D**) of cytoskeleton organization. Scale bars: 1 mm (**C**), 20 μm (**D**, first and third row) and 5 μm (**D**, second and fourth row).

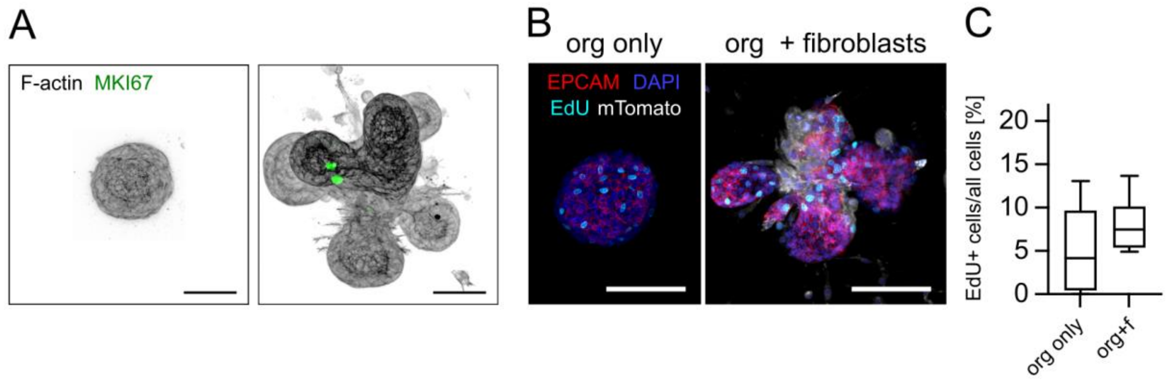


**Supplementary Figure 5. Proliferation in co-culture system. A**. F-actin (black) and MKI67 (green) staining in organoids cultured with fibroblasts, day 4 of culture. Scale bar: 50 μm. **B.** Photographs of organoid on day 4 of (co-)culture, EdU was administered 2 h pre-fixation. EPCAM (red), DAPI (blue), EdU (cyan), fibroblasts were isolated from *R26-mT/mG* mice (mTomato, white). Scale bar: 100 μm. **C.** Quantification of number of EdU+ cells of all cells in organoids in basal medium alone (org only) or with fibroblasts. The box and whiskers plot shows minimum, median, and maximum values, and second and third quartiles of data distribution. Statistical analysis: two-tailored *t*-test, not significant.

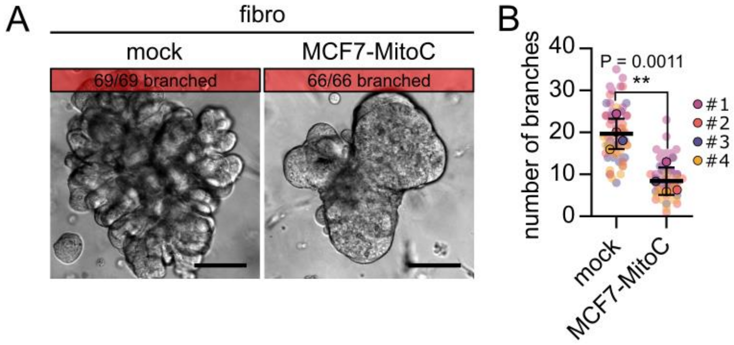


**Supplementary Figure 6. Spheroid proliferation is necessary for its branching in co-culture with fibroblasts. A.** Representative images of MCF7-ras spheroids in co-culture with fibroblasts on day 4 with spheroids formed from mock- or mitomycin C-treated MCF7-ras cells. The insets (top red bars) show proportion of branched spheroids out of all spheroids per condition. Scale bar: 100 µm. **B.** The plot shows number of spheroid branches/buds formed, with mean ± SD. Each lined dot represents mean of each experiment, each faint dot represents one spheroid, n = 4 independent experiments (coded by dot colors), N = 15-20 spheroids per experiment. Statistical analysis: two-tailored *t*-test.

### Supplementary Videos

**Supplementary Video 1. Mammary epithelial branching morphogenesis upon FGF2 treatment or fibroblast co-culture.** The video is composed of time-lapse videos capturing 5 days of epithelial morphogenesis in 3D organoid culture with no growth factor in the basal organoid medium (left), with FGF2 in the basal organoid medium (middle), or in fibroblast-organoid co-culture without addition of any growth factors to the basal organoid medium. Time is in hours. Snapshots from the videos are depicted in Fig. 1A.

**Supplementary Video 2. Fibroblasts dynamically interact with the epithelium.** Time-lapse video (combination of bright-field and fluorescence imaging) shows 2 days of epithelial morphogenesis in fibroblast (red)-organoid (white) co-culture (day 2-4).

**Supplementary Video 3. Fibroblasts form close contacts with epithelium in the organoid branching points.** 3D structure of organoid-fibroblast interaction. Single images are shown in Fig. 2C. Luminal cells (KRT8), red; basal cells (KRT5), blue; all cells (F-actin), white.

**Supplementary Video 4. *Myh9* knock-down in fibroblasts decreases their morphogenetic potential.** Time-lapse videos show 5 days of epithelial morphogenesis in co-culture with either control (left) or *Myh9* knocked-down fibroblasts (siRNA-mediated knockdown; si*Myh9*; right). Snapshots from the video are depicted in Fig. 3.

**Supplementary Video 5. *Myh9* knock-out in fibroblast decreases their morphogenetic potential.** Time-lapse video captures 5 days of epithelial morphogenesis in fibroblast-organoid co-culture with either control (Ad-GFP; left) or *Myh9* knocked-out fibroblasts (adeno-Cre-mediated knock-out; Ad-Cre-GFP; right). Snapshots from the video are depicted in Fig. 3.
